# Specific targeting of layer 4 for direct cortical stimulation is not necessary to induce reliable perception

**DOI:** 10.64898/2026.01.08.698338

**Authors:** Alexandre Tolboom, Guillaume Hucher, Daniel E. Shulz, Valérie Ego-Stengel

## Abstract

The primary somatosensory cortex (S1) processes tactile inputs through microcircuits involving connections between and within layers. In particular, layer 4 is the main first cortical target of tactile inputs. However, the need for the initial activation of layer 4 neurons in order to generate a tactile percept has not been studied yet. Here, we investigated whether optogenetic activation of layer 4 neurons first is necessary to induce perception, or whether stimulation of neurons in other layers may suffice. To achieve this, we generated mouse profiles expressing a light-sensitive opsin in excitatory neurons in different combinations of cortical layers, including or excluding layer 4. We generated these profiles by combining transgenic mouse lines and peripheral injections of PHP.eB adeno-associated viruses. After providing proof-of-concept for neuronal optogenetic control in vivo with PHP.eB viruses, we trained mice in a task involving the discrimination of moving optogenetic stimuli projected onto the whisker S1 area, thanks to patterned photostimulation. Mice learned to track the position of a light bar in order to obtain rewards. Direct activation of layer 4 was sufficient for the mice to learn the task, but not necessary. Indeed, activation of all layers but layer 4 induced learning. Moreover, the simultaneous direct activation of all layers accelerated learning. These findings contribute to a better understanding of the cortical mechanisms underlying somatosensory perception. They could also help to optimize artificial sensory stimuli to provide efficient feedback in cortical neuroprostheses.

## Introduction

In rodents, a single whisker twitch can trigger a cascade of neural activity — but how these spiking events lead to perception remains unclear. Indeed, the transformation of touch into a tactile percept is a complex process involving many actors and interplays in the central nervous system.

The primary somatosensory cortex (S1) is the dominant entry point of tactile inputs into the neocortex (Ferezou et al., 2007), integrating them in a behaviorally relevant manner (Pluta et al., 2017; Sachidhanandam et al., 2013; Sofroniew et al., 2015). S1 transient silencing or ablation impairs detection of tactile and vibro-tactile stimuli in rats and mice (Finger and Simons, 1976; Hong et al., 2018). During whisker-guided locomotion, broad inactivation of the S1 barrel cortex area (wS1), in which each barrel corresponds to one whisker (Petersen, 2019), also reduces wall-tracking accuracy in mice (Sofroniew et al., 2015). Furthermore, inhibiting S1 neurons during peripheral stimulation attenuates sensory responses as well as the animal’s readout of a given stimulation (Sofroniew et al., 2015). More generally, any perturbation of S1, whether activation or inhibition, can disrupt the integration of an ongoing tactile stimulus (Sun et al., 2021). Conversely, many studies have pointed out the sufficiency of S1 activation in the generation of tactile perception. For example, artificial stimulation of S1 in rodents can replace whisker touch during locomotion or tactile discrimination tasks (O’Connor et al., 2013; Sofroniew et al., 2015). In non-human primates, electrical stimulation of S1 produces behavioral responses and discrimination performance that are indistinguishable from those induced by natural skin stimulation (Romo et al., 1998). When implemented as an artificial tactile feedback in brain-machine interfaces, it can also guide motor control and virtual texture discrimination (O’Doherty et al., 2011). Finally, in humans, cortically-stimulated patients report tactile-like percepts, felt at consistent locations with respect to the somatotopic S1 map and including a wide variety of skin sensations (Flesher et al., 2016; Hughes et al., 2021; Lee et al., 2018; Tabot et al., 2015). Overall, these studies have demonstrated that artificial activation of S1 partially reconstitutes natural perception, especially by engaging the same neural networks (Pancholi et al., 2023).

Studies aiming to dissect neocortical circuits have highlighted a specific organization of the microcircuitry (Douglas and Martin, 2004; Harris and Shepherd, 2015; Petersen, 2019). Neocortical areas are organized in a laminar and columnar manner. Neuronal excitatory subtypes are distributed in specific proportions inside layers, and connect according to specific horizontal and vertical wiring diagrams, which differ slightly across areas and species (Harris and Shepherd, 2015). In rodent S1, stellate cells are the main neuronal subtype of layer 4 (L4) (Scala et al., 2019), and their activation generates the strongest excitatory influence on the cortical column (Lefort et al., 2009). The strongest intracortical ascending connections project from L4 to L2 and L3, while the main descending ones are exclusively targeting L5 and L6 (Lefort et al., 2009; Staiger and Petersen, 2021). Lemniscal inputs to S1 propagate through a core feed-forward canonical pathway - common to other primary sensory areas - in a given order (but see below): thalamus→L4→L2/3→L5→L6. This cortical circuit has an entry hub (L4) and two output layers projecting to subcortical and distant cortical structures (L5) or back to the thalamus (L6) (Kim et al., 2014; Olsen et al., 2012; Petersen, 2019; Petersen and Crochet, 2013).

However, other projections exist in parallel to this canonical pathway. First, regarding the entry of tactile information into the cortex, the posteromedial nucleus (POm) projects to L1 and L5a (El-Boustani et al., 2020), while the core of the ventral posteromedial nucleus (VPM) also targets the boundary between L5 and L6, in addition to targeting L4 of S1 (Feldmeyer, 2012; Petersen, 2019; Wimmer et al., 2010). Furthermore, strong functional projections have been observed laterally between cortical columns as well as vertically, and participate to intracortical computations (Moore and Nelson, 1998). For example, depending on stimulation parameters, activation of L4 of S1 can drive L5 pyramidal neurons (O’Connor et al., 2013) or suppress them by recruitment of fast-spiking inhibitory interneurons of L5 (Pluta et al., 2015). Spontaneous activity and sensory evoked activity strongly differ in their distribution of cortical layers, highlighting the complexity of the activity resulting from the patterns of connections (Beltramo et al., 2013; Peron et al., 2015; Sakata and Harris, 2009; Sofroniew et al., 2015; Zhao et al., 2016). Thus, although S1 is clearly involved in the generation of tactile perception, the underlying microcircuit mechanisms remain unclear. In particular, the importance of activation being initiated in L4 first, as in the canonical circuit, on perception and subsequent behavior, has not yet been studied.

Dissecting the circuits of sensory perception is now facilitated by genetic tools enabling excitation or inhibition of specific neuronal subtypes. Classical options to implement these methods are transgenic mouse lines and intracerebral injections of adeno-associated viruses (AAVs). The former option requires costly and time-consuming breeding schemes, which limit the implementation of new tools. The latter option provides strong neuronal transduction but requires invasive injections, and do not result in large and uniform transduction coverage. Indeed, intracerebral injections result in a small volume of tissue labelling with a strong gradient around the delivery site, a limiting factor for performing well-controlled large-scale imaging or photostimulation (Goertsen et al., 2022; Grødem et al., 2023; Leikvoll and Kara, 2023).

The AAV-PhP.eB serotype (Chan et al., 2017) delivered systemically is capable of crossing the blood-brain barrier. This cutting-edge method allows for long-term transgene expression in neurons after a single intravenous injection lasting a few minutes. Several studies have shown that the expression pattern in a given brain tissue is uniform, widespread, and stable over time. Moreover, many studies demonstrated the feasibility of 1- or 2-photon calcium imaging after PhP.eB-mediated GCaMP expression (Bharioke et al., 2022; Grødem et al., 2023; Michelson et al., 2019), even up to 8 months after injection (Leikvoll and Kara, 2023).

To our knowledge, no study has reported the use of blood-brain-barrier-permeable viruses to express light-activated proteins in the brain. Here, we used the PhP.eB serotype to produce widespread opsin expression for mesoscale activation of S1 neurons. Specifically, we aimed to express ChR2-H134R in subsets of cortical neurons by transfecting Emx1-Cre and Scnn1a-Cre mice. We obtained transgene expression in excitatory neurons of S1, in all layers except L4 for Emx1-Cre mice (Madisen et al., 2012) and in L4 only for Scnn1a-Cre mice (Kim et al., 2024). In addition to these two mouse lines, we also generated Emx1-Cre x Ai32 mice expressing the same opsin in excitatory neurons of all layers. We hypothesized that depending on the transgene expression pattern, particularly whether it included L4 or not, the same photostimulation on wS1 could lead to different detection and discrimination abilities. We tested this by training mice to discriminate the position of a dynamic photostimulation applied on wS1 thanks to mesoscale patterned optogenetics.

## Results

### PhP.eB-mediated transgene expression leads to heterogeneity across layers of the barrel cortex in Emx1-Cre mice

In order to test if the activation of S1 must include specific layers to generate perception, we generated mice with different profiles of opsin expression across layers of wS1. First, we injected a Cre-dependent viral preparation in the retro-orbital sinus bloodflow of 5 to 11-week-old Emx1-Cre mice (***Fig. 1A*)**. Viral genome contained a floxed gene construct coding for ChR2-H134R fused to eYFP. We chose the PhP.eB serotype for its ability to cross the blood-brain barrier and its tropism biased towards the central nervous system and neurons (Chan et al., 2017). After at least 4 weeks, we perfused the mice and imaged the fluorescence of brain sections. For all Emx1-Cre mice injected with the virus, histology revealed transgene expression with a large coverage of the whole neocortex, from the prefrontal areas to the visual areas (***Fig. 1B, Fig. Suppl. 1A)***. Consistent with previous studies using PhP.eB, we found regions of local higher fluorescence within structures: the CA2 region in the hippocampus (Grødem et al., 2023; Konno and Hirai, 2020; Mathiesen et al., 2020; Michelson et al., 2019) (***Fig. 1B, Fig. Suppl. 1A)***, the retro-splenial cortex in the neocortex (Michelson et al., 2019) (***Fig. 1B*),** and L5 in general among layers of the neocortex (Grødem et al., 2023; Michelson et al., 2019) (***Fig. 1B, Fig. Suppl. 1A)*.**

**Figure 1.**
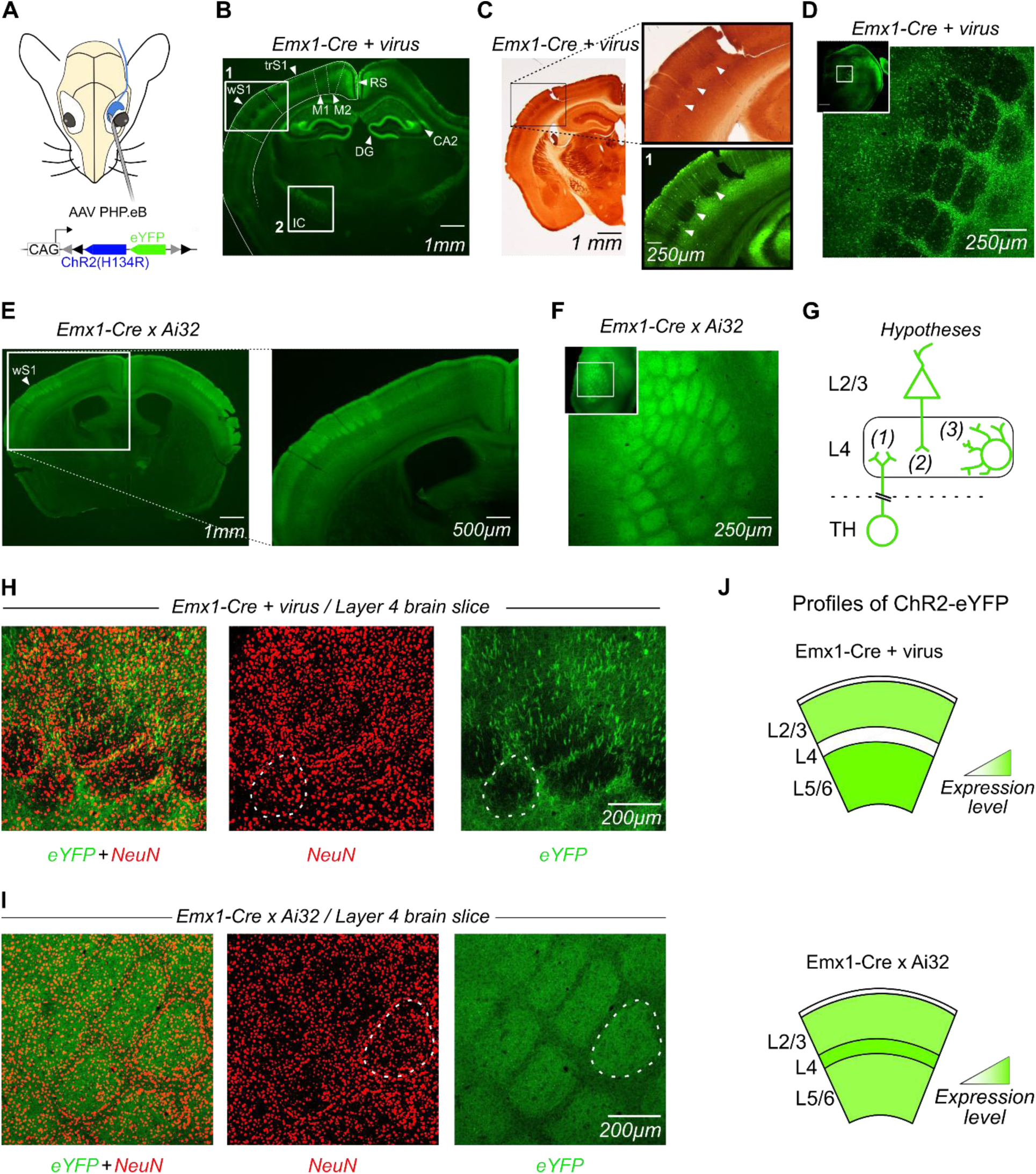
Opsin expression across cortical areas and across layers of the barrel cortex in transgenic and PHP.eB injected Emx1-Cre mice. **A Top,** Schematics of the intravenous injection. AAV PHP.eB particles were injected in the retro-orbital sinus of Emx1-Cre mice. **Bottom,** The viral genome contained insert gene coding for ChR2(H134R) fused to eYFP. Black and grey triangles indicate lox2272 and loxP sites. **B** Fluorescence image of a coronal section from an Emx1-Cre mouse brain showing expression of ChR2(H134R)-eYFP 12 weeks after AAV PHP.eB injection. The neocortex, hippocampus and white matter structures display high fluorescence. The whisker-associated primary somatosensory cortex (wS1) and the internal capsule (IC) are outlined in white boxes (see higher magnification in Fig. 1C & **Fig. Suppl. 1C**). RS: retro-splenial cortex; DG: dentate gyrus; CA2: subregion of the Cornus Ammonis (CA) in the hippocampus; M1: primary motor cortex; M2: secondary motor cortex; trS1: trunk region of S1. **C Left**, Section adjacent to the section displayed in panel **B** and treated with cytochrome C oxidase. **Top right**, Higher magnification image of wS1. The arrows show three barrels stained in layer 4. **Bottom right**, Corresponding fluorescence image of wS1 from the section displayed in panel B (white box 1). Arrows indicate low-density spots located in the regions corresponding to the barrels. Thick vertical fiber bundles cross all layers of wS1. **D Inset,** Fluorescence image of a tangential cortical section of an Emx1-Cre mouse brain 52 weeks after injection, centered on wS1 layer 4. Higher magnification shows low fluorescence in the barrels compared to septa. **E Left**, Fluorescence image of a coronal section of an Emx1-Cre x Ai32 mouse brain including the wS1 area. **Right**, Higher magnification shows high fluorescence in the barrels. **F** Same as D for an Emx1-Cre x Ai32 mouse brain. The higher magnification image shows high fluorescence in the barrels compared to septa. **G** Schematic of the three main hypotheses explaining the presence/absence of fluorescence in the neuropil of layer 4 in Emx1-Cre mice. The entities expressing or not the transgene could be: (1) thalamo-cortical neurons projections to L4; (2) projections of other neurons from the same column to the L4; (3) neurites of intra-barrel neurons. **H,I Left**, Transgene fluorescence (green) and NeuN immunostaining (red) from a tangential cortical section including layer 4 of wS1 from an Emx1-Cre mouse injected with PHP.eB mouse brain (**H**) and from an Emx1-Cre x Ai32 mouse (**I**). **Middle**, neuronal nuclei revealed by NeuN staining show a slightly higher concentration of somas at the barrel walls. Z-projection of 5 scan sections representing a total thickness of 20 µm. An example barrel is outlined in white. **Right**, Z-projection of transgene eYFP fluorescence from the same sections. wS1 barrels display low fluorescence in Emx1-Cre mouse injected with PHP.eB (**H**) and high fluorescence in Emx1-Cre x Ai32 mouse (**I**). Note that in panel F, the sparse bright eYFP spots do not colocalize with somatic NeuN spots, suggesting they are due to neuropil bundles (see also **Fig. Suppl. 1**). **J** Schematic representation of the estimated transgene expression level in the membranes of neurites and somata across cortical layers using the PHP.eB viral strategy (**Top**) and the transgenic mouse line strategy (**Bottom**).

In these Emx1-Cre + virus mice, given the promoter, we anticipated that the targeting of all cortical excitatory neurons would result in a homogeneous fluorescence profile across wS1 layers. Surprisingly, we observed fluorescence-free spots at a middle depth of wS1 area (white rectangle 1, ***Fig. 1B-C***). Thick fluorescent fibers crossed these dark spots vertically, probably corresponding to bundles of apical dendrites from deep layers neurons. Given their size of around 300 µm, their laminar depth and their overall distribution, we hypothesized that the dark spots corresponded to the barrels. To test this, we cut 2 brains in thin 50-µm slices and treated every other slice with cytochrome C oxydase to reveal the barrels in the barrel field of S1. This enabled a comparison of the eYFP fluorescence patterns against the classical barrel staining in pairs of neighboring slices. Fluorescence-free areas were found to systematically match labelled barrels (white arrows in ***Fig. 1C*)**. To confirm these observations, we also imaged tangential slices of the cortex (***Fig. 1D*)**. Fluorescence-free oval spots were observed in row and arc-like formations, validating that we were visualizing L4 barrels of wS1. Sparse bright dots, most probably due to the putative bundles of apical dendrites already noticed in coronal sections, were scattered throughout barrels and septa. These findings confirm a diminished expression of the transgene in L4 neuropil of wS1 relative to other layers, particularly inside barrels.

To further investigate the absence of expression in the barrels, we examined the expression of the same construct (ChR2-H134R fused to eYFP) under the same Emx1 promoter, but this time expressed by a transgenic mouse line crossing strategy (Emx1-Cre x Ai32 mice). The transgene expression was robust in all cortical layers, including L4 barrels, which were even slightly brighter than neighboring layers (***Fig. 1E-F, Fig. Suppl. 1E)***. Thus, we obtained two different expression patterns in these two mouse groups: all layers labelled in crossed transgenic mice Emx1-Cre x Ai32, and all layers except L4 in Emx1-Cre mice injected with a PHP.eB virus.

We explored several hypotheses that could explain this striking difference of the fluorescent protein level in L4 barrels. First, comparison of eYFP and NeuN patterns showed that eYFP fluorescence did not co-localize with nuclei of L4 neurons in either mice group (***Fig. Suppl. 1G-H***). Indeed, this is in line with the expected localization of the ChR2-eYFP fused protein at the neuronal membrane, mostly in dendrites and axons. This is also consistent with the strong expression levels in fiber tracts such as the anterior and internal commissures (***Fig. 1B, Fig. Suppl. 1A-C)***, cerebellar peduncles (***Fig. Suppl. 1A***) and pyramids of the medulla (***Fig. Suppl. 1D***), as well as the strong expression in regions of the dentate gyrus that do not contain the cell bodies revealed with DAPI staining (***Fig. Suppl. 1B***). Overall, we conclude to no or very little localization of the opsin in cell bodies in general, and rather that the fluorescence originates from the neuropil (dendrites, axons).

Based on the known constituents of the neuropil inside barrels (Staiger and Petersen, 2021), the higher fluorescence in Emx1 x Ai32 mice could be due to a higher fluorescence in (1) axons from thalamocortical cells, (2) dendrites and axons of wS1 excitatory neurons with somas in more superficial and deeper layers, and/or (3) dendrites and axons of local L4 excitatory neurons ***Fig. 1G***. We first considered whether the thalamocortical axons and synapses, known to be abundant in barrels (Egger et al., 2008; Oberlaender et al., 2012; Woolsey and Van der Loos, 1970) could contribute to the fluorescence. However, thalamic neurons of Emx1-Cre mice do not express the Cre recombinase (Gorski et al., 2002) (see also Discussion), ruling out this possibility. Second, we considered whether dendrites and axons of cortical neurons from upper and lower layers could explain the concentration of fluorescence in barrels relative to septa in Emx1 x Ai32 mice. Given that extra-barrel cortical neurons project very few dendrites and axons in barrels (Lefort et al., 2009; Staiger and Petersen, 2021), except thick apical trunks that we could indeed observe as vertical bundles, this is also very unlikely. Thus, we turned to our third hypothesis for the source of intra-barrel fluorescence, the intra-barrel excitatory neurons (***Fig. 1G*)**. In both types of mice, we labelled the somata of neurons with anti-NeuN immunostaining in tangential sections in L4, revealing a relatively homogeneous distribution of somas in barrels and a slightly higher concentration in septa (***Fig. 1H-I***). Most of these cells are putatively excitatory stellate cells (Narayanan et al., 2017; Scala et al., 2019; Staiger and Petersen, 2021), known to have a strong dendritic and axonal arborization within the barrel extending away from the septa (Oberlaender et al., 2012; Staiger and Petersen, 2021). Thus, we conclude that the observed difference in fluorescence patterns is due to transgene presence in the membrane of neurites and soma of excitatory neurons in L4 barrels in Emx1-Cre x Ai32 mice, and not in Emx1-Cre mice injected with the virus (***Fig. 1J***, see Discussion).

As already mentioned, the use of a membrane-bound reporter (ChR2-H134R fused to eYFP) created a global fluorescence of the neuropil, making it difficult to identify the cell bodies of neurons expressing the transgene. To investigate this further, a cytoplasmic reporter was used (GCaMP8m) in two pilot animals. Emx1-Cre mice were injected in the retro-orbital sinus with AAVs of PhP.eB serotype containing a floxed GCaMP8m gene under CAG promoter (same construct as for ChR2-H134R). We imaged coronal sections of the brain and found results consistent with those of mice injected with ChR2-H134R construct (***Fig. Suppl. 2)***. Fluorescent somata were found in L2/3 and L5, while very few positive somata were observed in L1 and L4. This result confirms that the expression bias away from L4 excitatory neurons is probably due to the PHP.eB serotype in Emx1-Cre mice, and not to the particular transgene cargo expressed (see Discussion).

### Transgenic and PhP.eB viral strategies both lead to transgene expression in layer 4 of wS1 in Scnn1a-Cre mice

To test the necessity and sufficiency of activating L4 of wS1 first in order to drive sensory perception, we then aimed to express the opsin transgene only in L4. To do so, we performed the same viral injections in Scnn1a-Cre mice, a transgenic line specifically targeting L4. As expected, confocal images revealed high expression in L4 of wS1 (***Fig. 2A***). We noticed highly fluorescent barrel-like shapes organized side by side. This was confirmed by tangential section images showing barrels filled with fluorescence, and arranged according to the somatotopic map (***Fig. 2B***). In addition, L4 of other sensory cortices like the posterior parietal cortex area (PPC), the primary auditory cortex and dorsal associated areas (A1, AUD) also showed high expression (***Fig. Suppl. 1F***). All these observations are in line with the Scnn1a gene expression profile described in the literature (Kim et al., 2024). Following the same protocol as for Emx1-Cre mice, we also imaged brain slices from Scnn1a-Cre x Ai32 mice. Apart from an even higher fluorescence signal in L4 compared to other layers, the spatial pattern of expression was similar to the pattern of Scnn1a-Cre mice injected with the PHP.eB virus (***Fig. 2CD*).**

**Figure 2.**
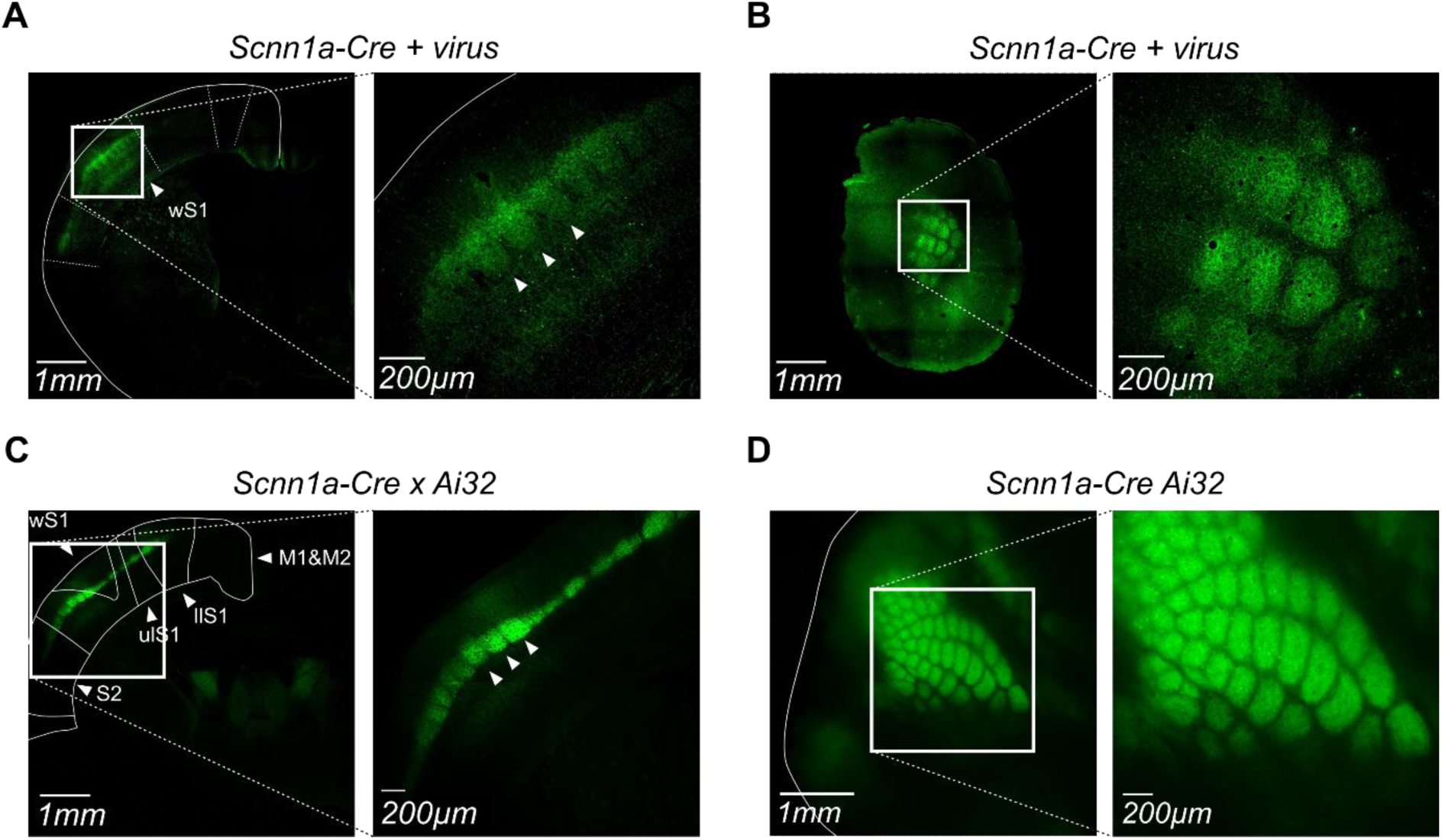
Similar transgene expression profiles in the barrel cortex of Scnn1a-Cre mice following injection of AAV PHP.eB particles or crossing with a conditional transgenic mouse line. **A Left** Fluorescence image of a coronal section from a Scnn1a-Cre mouse brain 7 weeks after AAV PHP.eB injection, centered on wS1 (white box). **Right** Higher magnification image of wS1. White arrows indicate layer 4 barrels. **B Left** Fluorescence image of a tangential section from another Scnn1a-Cre mouse 7 weeks after injection. **Right** Higher magnification image of wS1. **C** Same as **A** from a Scnn1a-Cre x Ai32 mouse brain. Only layer 4 barrels are labelled in wS1. llS1: lower limb S1 area; ulS1: upper limb S1 area; S2: secondary somatosensory area. **D** Same as **C** for a tangential section from a Scnn1a-Cre x Ai32 mouse brain, centered on wS1 (white box).

Thus, we generated three laminar profiles of ChR2-H134R expression in excitatory neurons in S1: “all layers” (Emx1-Cre x Ai32), “all layers except L4” (Emx1-Cre + virus) and “only L4” (Scnn1a-Cre + virus). Note that inhibitory neurons of all layers, which include all L1 neurons, are not activated by our photostimulation.

### Photostimulation of layer 4 of wS1 induces neuronal activation that can be detected and reported by mice expressing ChR2-H134R after AAV-PHP.eB injection

To verify that the opsin expression was sufficient to trigger action potentials after delivery of the transgene using the PHP.eB serotype, we performed electrophysiological recordings in vivo while stimulating the cortical surface with blue light.

A silicon probe was inserted obliquely through a small opening of the glass cranial window (see Methods) in the wS1 cortical area of an anaesthetized Emx1-Cre mouse injected with AAV PHP.eB (***Fig. 3A* *Left)***. We projected a series of light spots, spatially organized in a grid on the cortical surface, using a Digital Light Processor (DLP) (***Fig. 3A* *Right)***. A new spot was projected every 100 ms in a pseudo-random order, and consisted of 5 flashes of 5 ms each and separated by 5 ms (***Fig. 3C, Fig. Suppl. 3B)***. After spike sorting and curation (see *Methods for validation criteria*), we obtained 5 single units (example spike shape and autocorrelogram in ***Fig. Suppl. 3C).*** For all recorded units, the mean firing rate increased during the stimulation period compared to the baseline (Wilcoxon test, p = 0.0312, ***Fig. 3B*).** The response time course of the example unit to spot 1 and spot 11 show no spiking after the projection of spot 1 (***Fig. 3C*, Left**) while a phase-locked activity was induced by the flashes of spot 11 (**Right**). Indeed, spots 10 and 11 evoked a large increase in mean evoked activity compared to baseline, while other spots did not evoke activation (***Fig. 3D*)**. Similar patterns of responses were obtained for 4 of the 5 recorded units, for which spots 7, 8,10 and 11 evoked the highest firing rate (***Fig. Suppl. 3D***). We conclude that PHP.eB-mediated opsin expression can be used to activate neurons in the neocortex even with a brief stimulus (5 ms). The temporal and spatial characteristics of these responses, including long delays in some cases **(*Fig. 3C*)** are compatible with previous recordings in the lab in Emx1-Cre x Ai32 and Emx1-Cre x Ai27 mice, as well as responses to optogenetic flashes in *in vivo* published data (Ceballo et al., 2019b).

**Figure 3.**
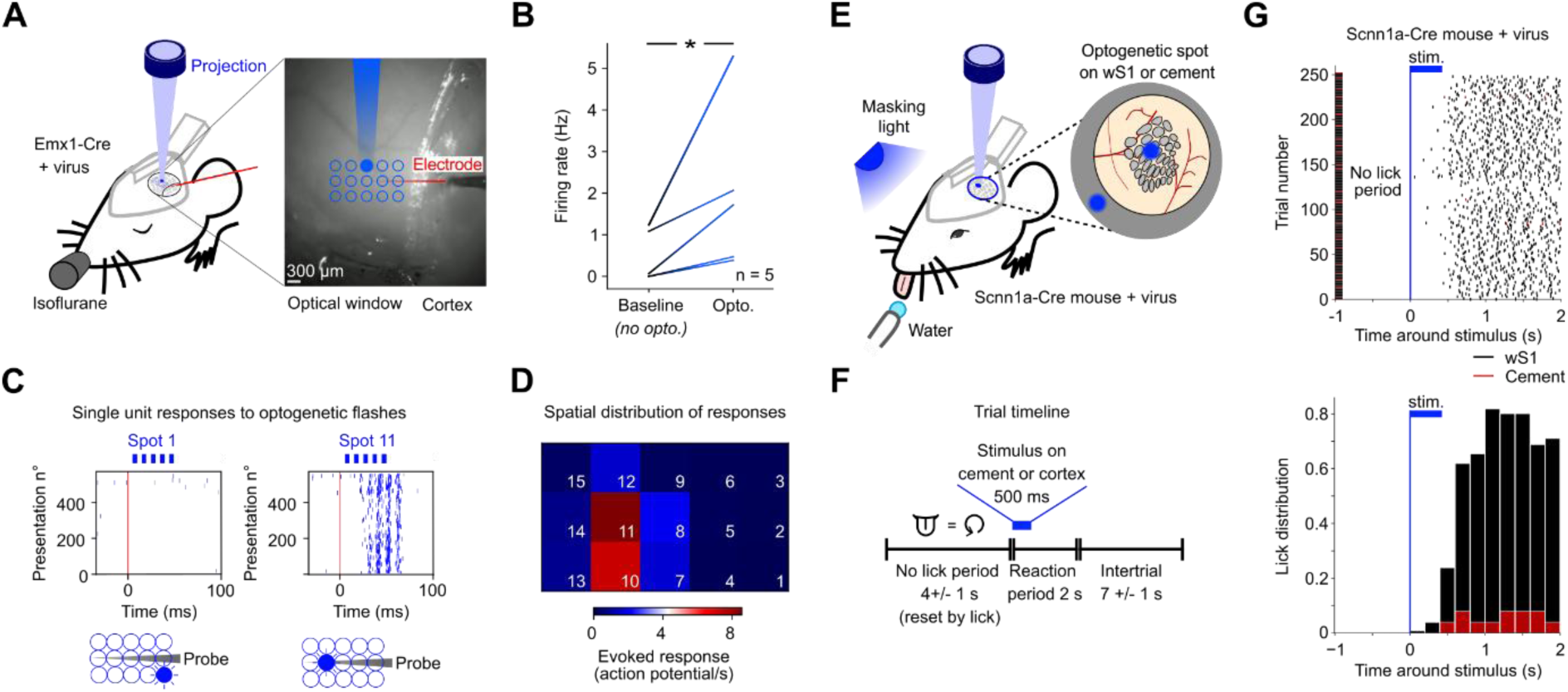
Photostimulation of wS1 activates cortical neurons and creates detectable percepts in mice expressing ChR2-H134R after AAV PHP.eB injection. **A Left**, Schematics of the setup. An Emx1-Cre mouse injected with AAV PhP.eB particles was head-fixed under anaesthesia. A 32-channel silicon probe was inserted below the cranial window through a small opening. We recorded neuronal activity while projecting light spots with a digital projector (DLP) through the intact part of the window. Fifteen different spots were projected in pseudo-random sequences, according to a 3×5 stimulation matrix centered above the inserted probe. **Right**, image of the cranial window with the electrode penetrating the cortex from the side. A sketch of the light spots and of the electrode shank are overlaid. **B** Mean firing rates of all 5 single units before and during the photostimulation protocol (not specifically after a given spot projection). All units showed an increased activity (Wilcoxon test, * p < 0.05). **C** Raster plots of action potentials for one single unit, centered on the beginning of the first flash of spot 1 (Top) and spot 11 (Bottom). Four to five phase-locked peaks of response were observed in response to spot 11 while spot 1 didn’t trigger any activity. **D** Average increase in the number of action potentials per second during the 80 ms following a light spot onset compared to the baseline period activity, for each of the 15 light spots. Same neuron as panel C. **E** Detection task. Scnn1a-Cre mice injected with AAV.PHP.eB (carrying the gene coding for ChR2-H134R) or control mice were head-fixated under the same DLP system as in **A.** Mice were awake and water-restricted. For each trial, an optogenetic spot (300 µm diameter) was projected onto the window, either targeting the C2 barrel or targeting the dental cement (sham catch trial). The mouse had to lick a spout to obtain a water reward in the two seconds following the stimulus start. A masking blue light was directed towards the eyes to mask the projection of spots. **F** Timeline of a single trial. Each trial started with a No Lick Period of random duration (between 3 and 5 s). This period was reset by each lick to force the mice to stop licking and focus on detecting the stimulation. Then, the stimulus was projected during 500 ms: 50 flashes of 5 ms separated by 5 ms (same duty cycle as in Fig. 3C). The trials in which the spot was projected onto the cement accounted for 10% of the total, with one catch trial randomly placed per group of 10 trials. From the first flash, the mouse had 2 seconds to lick the spout twice in order to receive a reward (Reaction period). Licks to both cortical and cement spots were rewarded by a water droplet. Finally, an intertrial period (6 to 8 s long) allowed the mouse to drink its reward before the start of the next trial. **G** Licking activity for the Scnn1a-Cre mouse injected with the virus (“only L4”) during session 9 of the detection task. **Top**, Raster plots of licks during all trials of the session. The type of trial (spot onto cortex or cement) is shown by a colored tick on the left (cortex black, cement red) and by the matching color of the licks. **Bottom**, Mean lick distribution computed over all trials of the session, for trials in which the light spot was projected onto the cortex (black) or cement (red). Time 0 corresponds to the beginning of the first optogenetic flash.

Having validated that photostimulation can drive spiking activity in an anesthetized animal, we tested whether photostimulation can lead to perception in the awake state. We implemented a behavioral task under head-fixation and water restriction. Each trial started with a ‘No Lick Period’ (3 to 5 s) followed by the presentation of a light spot of 300 µm diameter (similar to spots in ***Fig. 3A***) projected during 500 ms onto the C2 barrel or, as a control, onto the dental cement surrounding the cranial window (***Fig. 3EF*)**. A masking blue light was directed toward the mouse’s face to limit visual detection of the spots. The mouse had two seconds to react to the spot by licking a spout, in order to obtain a water reward. Both stimuli, onto the cortex and the cement, were rewarded (***Fig. 3F*)**. This ensured that if the mouse could detect the light visually by reflections in the setup, rather than through neuronal activation, it should lick similarly for both spots, an outcome that should be obvious on lick raster plots (***Fig. Suppl. 4A Bottom***).

We trained a Scnn1a-Cre mouse injected with AAV PHP.eB in this task. We computed the raster plots of licks and the resulting lick distributions centered on the start of the stimulation for the cortical and cement spots (***Fig. 3G*)**. After 9 sessions of training, the mouse initiated bursts of licks after almost all optogenetic spots projected onto wS1 (95%), while mostly ignoring the ‘cement’ spots (8%). This difference indicated that the optogenetic detection task was not solved by seeing light patterns reflected in the setup. We reached the same conclusion in previous work in transgenic mice expressing channelrhodopsin across all layers (Abbasi et al., 2018).

To further exclude that detection of the spot could be achieved by light propagation from the cortical surface to the retina through the brain tissue, we trained two control Scnn1a-Cre mice that had not been injected with PHP.eB in the same task, and found that these mice could not detect the photostimulation spots (***Fig. Suppl. 4)***. Overall, we conclude that mice expressing channelrhodopsin with the PhP.eB viral strategy can detect and report the presence of a static optogenetic light spot thanks to direct activation of neurons in wS1.

### Activation of layer 4 first is sufficient, but not necessary, to discriminate spatiotemporal optogenetic patterns projected onto wS1

In contrast to the static spot stimulation used above, somatosensory stimuli are intrinsically dynamic. For example, when a mouse is whisking against an object, it generates global patterns of fast deflections across arcs and rows of whiskers. Therefore, to test the discrimination abilities of mice, we turned to cortical stimulation patterns exhibiting more complex spatial and temporal dynamics. Specifically, we compared the performance of the three groups of mice (“all layers except L4”, “only L4” and “all layers”) in a task requiring to discriminate the position of a continuously moving cortical optogenetic stimulus. For the “all layers” group, we used data previously acquired in the laboratory obtained with transgenic mice identical to the Emx1-Cre x Ai32, except for the reporter fused to the opsin, tdTomato instead of YFP (Emx1-Cre x Ai27, (Lassagne et al., 2022)).

Head-fixed mice had to report the position of a dynamic blue light bar projected on wS1 through a glass cranial window of 5 mm diameter **(*Fig. 4A*)**. The light bar was set to rotate in a disk around the C2 barrel (***Fig. 4B*).** In order to position the disk accurately, we performed intrinsic optical imaging after the cranial window surgery while stimulating single whiskers to map at least 3 functional barrel areas, including the C2 barrel (***Fig. 4C*)**. During trials, the light bar rotated smoothly across the barrel cortex, gradually activating neighboring areas. At the start of each trial, the bar appeared in a caudal position, then moved in either direction with various dynamics (***Fig. 4D, Fig. Suppl. 5)***. The rotation space was split in three zones with different behavioral consequences. A lick when the bar was in the No-lick zone aborted the trial and initiated a 5-s long intertrial. On the contrary, the mouse received water rewards after licking when the light was in the Rewardable zone. Licks while the bar was in the Neutral zones had no effect. It should be noted that if the bar entered the Rewardable zone and the mouse did not lick, then no reward was delivered (details in (Lassagne et al., 2022)).

**Figure 4.**
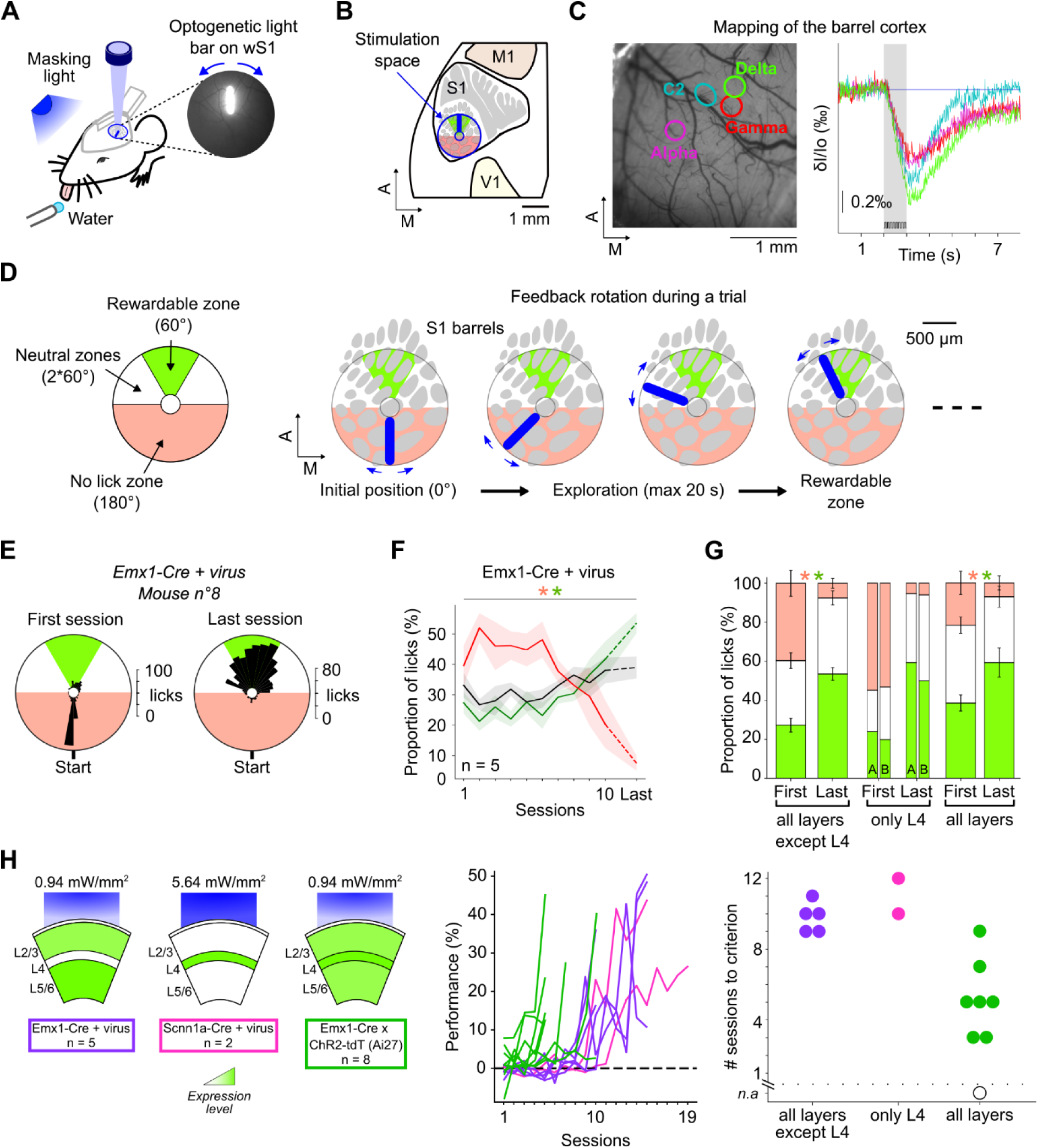
Discrimination of a dynamic cortical stimulation over wS1 is possible with direct activation of only L4, but also with other activation strategies. **A Left,** Sensory-guided licking task. A rotating light bar was projected on wS1 through a cranial window by a digital light processor (DLP). A water tube detected licks and delivered rewards when adequate. **Right,** An example image of the cortical surface during optogenetic stimulation (the white bar corresponds to blue light in grey scale). **B** Position of the stimulation space (blue circle) on the cortical functional map (grey) in S1. The space is centered on the C2 barrel and is restricted to the wS1 area. The map contours were adapted from (Knutsen et al., 2016), and (Vanni et al., 2017), and positioned according to intrinsic imaging of several barrels (panel C). **C** Example mapping of wS1 barrels under anesthesia prior to training. **Left,** Contours of intrinsic imaging responses to single whisker stimulations (Alpha, Gamma, C2 and Delta whiskers) superimposed on an image of the cortical surface. **Right**, Temporal light fluctuations (ΔI/I0) of the corresponding regions of interest during trials. The stimulation period is indicated by the grey shading. **D Left,** Different zones of the stimulation space. When the bar was in the Rewardable zone (green), licks were rewarded. In the No-lick zone (red), a lick ended the trial. In the Neutral zones (white), licks were ignored. **Right,** Snapshots of an example trial showing the displacement of the light bar on the barrel cortex. **E** Spatial distribution of the optogenetic bar angle at all lick timestamps for one example Emx1-Cre mouse trained 5 weeks after AAV PHP.eB injection, for the 1st and the 10th (Last) sessions. **F** Average proportions (mean ± s.e.m) of all licks for which the optogenetic bar was in the Rewardable (green), No-lick (red), and Neutral zones (black) during training of Emx1-Cre mice injected with PHP.eB of the “all layers except L4” group (n = 5). The “Last” session consists of the 10th session for two mice and the 15th for three mice. Wilcoxon test, * p < 0.05. **G** Average proportions (mean ± s.e.m) of all licks during training (same quantification as **F**). **Left,** Mean values from the first and last sessions for Emx1-Cre mice injected with PHP.eB (n = 5). **Middle,** Same for two Scnn1a-Cre mice (A and B) injected with PHP.eB (n = 2). **Right,** Same for Emx1-Cre x ChR2-tdT mice (n = 8). The “Last” session differs between groups, see panel H. Wilcoxon test * p < 0.05. **H Left,** Schematics of expression profile for the three groups: ‘all layers except L4’ Emx1-Cre + virus (purple), ‘only L4’ Scnn1a-Cre + virus (pink) and ‘all layers’ Emx1-Cre x Ai27 (green) mice. Blue light intensity was adapted to the cortical depth of opsin-expressing cells (see Methods). **Right,** Individual learning curves, quantified by the percentage of rewarded trials normalized by subtracting the chance level (see Methods). Same color code. **I** Learning speed measurement. Number of training sessions (one per day) to reach at least 10% of rewarded trials after normalization (corresponding to around 30% of raw percentage, **Fig. Suppl. 6**). Each circle represents a mouse. Same color code as panel **H**. The empty circle (n.a) represents an animal that never exceeded the chance level.

We trained five Emx1-Cre and two Scnn1a-Cre mice injected with PHP.eB particles (carrying the gene insert coding for ChR2(H134R), same as before) in this task. Since the only photoactivatable cells are located deeper in the cortex in Scnn1a-Cre mice, we adjusted the light intensity accordingly (***Fig. 4H* *Left,*** see *Methods*). During the first sessions, the mice licked frequently, often aborting trials. Most of the licks occurred at the very beginning of the trials near the Start position, probably in reaction to the appearance of the optogenetic bar (***Fig. 4E* *Left,*** representative example Emx1-Cre + virus mouse). After training, mice refrained from licking while the bar was in the No-lick zone and started lick bursts while close to the target Rewardable zone (***Fig. 4E* *Right).*** Population analysis on the five Emx1-Cre mice injected with PhP.eB (“all layers except L4”) confirmed this redistribution of lick angular positions: across sessions, mice significantly increased the percentage of licks while the bar was in the Rewardable zone, and decreased the percentage of licks for the No-lick zone (Wilcoxon tests, p = 0.0312 for licks in green and red zones, ***Fig. 4F*)**. Similar redistributions of licks were observed on the two Scnn1a-Cre mice injected (“only L4”) and the Emx1-Cre x Ai27 mice (“all layers”) (***Fig. 4G***). We conclude from this analysis of lick distributions that mice are able to report the angular position of the bar stimulus in all groups of animals, whether expressing the opsin in all layers, all layers except L4 or L4 only.

To further quantify learning across training days, we computed the percentage of rewarded trials per session. In the three groups, mice increased this percentage with training, although some of the mice needed many more sessions than others to learn the task (***Fig. Suppl. 6***). One potential caveat to this quantification is that mice could have just learnt the average time between the trial start and the entry in the Rewardable zone (around 10 seconds, ***Fig. Suppl. 5B***), which would allow them to perform relatively well without actually discriminating the position of the stimulation bar. To rule out this temporal strategy, we computed a chance level for each session by shuffling licking patterns and trial trajectories (see *Methods* and (Lassagne et al., 2022)). By subtracting this chance level, we obtained a normalized percentage of rewarded trials for each session (***Fig. 4H*)**. With both raw and normalized metrics, we noted that mice from the “all layers” group learned the task much more quickly than the others (***Fig. 4H*)**. In the “all layers except L4” and “only L4” groups, mice performed at chance level during the first 8 to 10 days (***Fig. 4H* *Right, Fig. Suppl. 6AB***). Learning appeared around day 9 to 12. For the last session of training, the normalized percentage of rewarded trials reached on average 33% for the “all layers except L4” mice (days 10^th^ to 15^th^ n = 5, Wilcoxon test, first vs. last session, p = 0.0312) and 35% for the “only L4” mice (days 15^th^ and 19^th^, n = 2) (***Fig. Suppl. 6AB)***. By contrast, in the “all layers” group, most mice learned the task quickly during the first 5 days of training (***Fig. 4H* *Right, Fig.Suppl. 6C***). Their performance in the last session after only 5 to 10 days reached 22% (n = 8, Wilcoxon test, first vs. last session, p = 0.0039). This difference in learning speed was quantified by the number of sessions it took each mouse to reach a criterion of 10% normalized rewarded trials (approximately 30% raw rewarded trials) **(*Fig. 4I*)**. “All layers” mice reached the threshold in 5.3 days, against 9.8 days for “All layers except L4” and 11 days for “only L4” mice (***Fig. 4I*).**

Overall, these results show that mice can perceive and discriminate cortical activation no matter in which layer it is initiated, but unexpectedly, mice learn faster when all layers are synchronously activated.

## Methods

### Animals and preparations

All animal experiments were performed according to European and French law as well as CNRS guidelines and were approved by the French ministry for research (Ethical Committee 59, authorization #25932). The data were obtained from 12 adult (5 to 11 weeks old) Emx1-Cre mice (Jax strain #005628), 6 adult Scnn1a-Tg3-Cre mice (Jax strain #009613), 4 adult Emx1-Cre x Ai32 mice (Ai32: Jax strain #024109, (Madisen et al., 2012)), 2 adult Scnn1a-Cre x Ai32 mice and 8 adult Emx1-Cre x Ai27 mice (Ai27: Jax strain #012567, (Madisen et al., 2012)).

We injected AAV PHP.eB particles (Addgene #127090) to obtain a Cre-dependent expression of ChR2-H134R fused to eYFP in excitatory neurons in all cortical layers (10 Emx1-Cre mice) or in L4 (4 Scnn1a-Cre mice). These particles have been designed to cross the blood-brain barrier and therefore infect neurons after intravenous injection (Chan et al., 2017). We diluted 2.1 ×10^11^ viral genomes in saline to obtain 100 µl of solution in a 1-mL insulin needle. Two additional Emx1-Cre mice were injected with AAV PHP.eB particles (Neurophotonics #1822-aavphp-eb) to obtain a Cre-dependent expression of GCamP8m (4.0 ×10^11^ viral genomes in saline, 100 µl of solution). We injected the right retro-orbital sinus of mice immediately after a 2 to 3 minutes isoflurane anaesthesia induction or during a Ketamine anaesthesia. We ensured there was no backflow during the injection. For the ketamine anaesthesia, we performed intra-peritoneal injections of a mix of Ketamine (60 mg/kg) and Medetomidine (1 mg/kg), and sub-cutaneous injections of atipamezole (2 mg/kg) to reverse anaesthesia. All injected and perfused mice later showed a positive histology.

Two weeks later, a cranial window surgery was performed under isoflurane anaesthesia in 100% air. In total, we implanted 5 virus-injected Emx1-Cre mice, 2 virus-injected Scnn1a-Cre mice, 2 control Scnn1a-Cre mice and 8 Emx1-Cre x Ai27 mice. Isoflurane concentration was adjusted in the range of 1 to 4% depending on mouse state, assessed by breathing rate and response to tail pinch. The scalp was resected after local anaesthesia induced by lidocaine (200 mg/L, 0.1 mL) and conjunctive tissues were removed. A head-post was glued (cyanoacrylate glue) to the skull, then strengthened with dental cement. A 5-mm diameter craniotomy was then performed while preserving the dura, centered on the stereotaxic coordinates of the C2 barrel in the left primary somatosensory cortex (P1.5-L3.3 mm). A glass optical window of diameter 5 mm was glued to the borders of the craniotomy. The remaining exposed skull was covered with dental cement. At the end of the surgery, we administered subcutaneously an analgesic in the back (2 mg/mL meloxicam, 0.1 mL). Ten days later, the clarity of the optical window was assessed, and if adequate, intrinsic imaging was performed to locate the S1 barrels (see below).

### Intrinsic imaging

Intrinsic imaging was performed under light isoflurane anaesthesia (0.9 to 1.4%). During an imaging session, a single whisker was stimulated with a piezoelectric bender (Physics Instruments, 100 Hz, 5 ms square wave deflection during 1 s) while red light (625 nm) was projected on the window. A CCD camera acquired 659*494 pixels images at 60 Hz. The images were analyzed for space-time fluctuations in luminescence (Optimage, Thomas Deneux, NeuroPSI). These intrinsic imaging sessions were used to locate the C2, Alpha, Gamma and Delta barrels. Details are available in (Abbasi et al., 2018).

### Histology

Several weeks after the injection or at the end of the behavioral experiments, mice were deeply anesthetized with 5% isoflurane, euthanized by an intraperitoneal injection of pentobarbital (730 mg/kg) and transcardially perfused with phosphate-buffered saline (PBS) followed by paraformaldehyde (PFA). For Emx1-Cre mice, we perfused the animals 4 to 12 weeks after viral injection (n = 6) and 52 weeks after (n = 2). Scnn1a-Cre mice were perfused 6 weeks after injections (n = 2). One Emx1-Cre mouse injected with the virus carrying the GCamP8m gene was not perfused, explaining the high background fluorescence in ***Fig. Suppl. 2*** (n°13, right image). All the brains were immersed in a 4% PFA solution overnight and then stored in PBS for two days before being cut. We collected coronal slices on the whole brain (80 µm thick) or tangential slices centered on wS1 (100 µm) using a vibratome (Leica VT1000S).

To measure fluorescence in the wS1 barrels, we cut thinner coronal slices (50 µm) and stained one out of two with cytochrome C oxidase. Then, we imaged the staining of the barrels on half of the slices and the fluorescence signals on the other half. Finally, we manually realigned pairs of adjacent slices to superimpose the two images. To confirm our results, we also imaged fluorescence signals directly on cytochrome C oxidase-stained slices. The results were identical. This protocol was applied for 2 Emx1-Cre mice injected, 2 Emx1-Cre x Ai32 mice and 2 Scnn1a-Cre x Ai32 mice. For all the mice, barrels estimated with eYFP signals matched the cytochrome stainings. Images were acquired using a Nikon Eclipse 90i (epifluorescence and cytochrome C oxidase imaging) and a Zeiss LSM880 Confocal AiryScan (objectives Plan-Apochromat 10x/0.45 M27 and Plan-Apochromat 20x/0.8 M27).

In order to estimate somata spatial distribution in the barrels, we performed NeuN immunostaining on tangential slices of L4 of wS1. We stained slices with a primary chicken anti-NeuN antibody (ThermoFisher #PA5-143567, dilution 1:450) followed by an Alexa Fluor 594 secondary antibody (ThermoFischer #A-11042, dilution 1:200). Confocal images were acquired with a Zeiss LSM880 Confocal AiryScan.

### Electrophysiological recording of optogenetically-induced activity

To measure spiking activity *in vivo*, we conducted an extracellular electrophysiological recording in an Emx1-Cre mouse 8 weeks after viral injection, under anaesthesia (1.5% isoflurane). A probe (32 channels, 1 shank of A2×32-5mm-25-200-177, Neuronexus) was inserted obliquely through a small opening of the wS1 cranial window while projecting light spots through the intact part of the window. We performed these optogenetic stimulations using a Digital Light Processing module (DLP, Vialux V-7001, 462 nm blue LED) to ensure precise patterns of light. Electrophysiological signals were recorded (Blackrock acquisition system) during a baseline period, followed by the optogenetic stimulations period. The light patterns consisted of single spots of 250 µm diameter, from a 3 x 5 matrix of spots centered on the probe insertion site. Spots were delivered every 100 ms in a pseudo-random order. Each spot consisted in 5 flashes of 5 ms, separated by 5 ms (50% duty cycle – see ***Fig. Suppl. 3B***).

### Spike sorting and analysis of single-unit activity

We analyzed the electrophysiological data using the SpikeInterface software and SpykingCircus2 sorting algorithm. We applied a bandpass filter (300-9000 Hz, Bessel filter, 2^nd^ order) and a common median reference. Sorted units were automatically merged based on correlogram and waveform template similarity, estimated distance between units and merging quality score. After applying curation criteria (inter-spike-interval violation ratio < 0.1, amplitude cut-off < 0.1) we manually validated units using Phy2.

### Detection task of an optogenetic spot projected over wS1

Three Scnn1a-Cre mice (one injected with viral particles and two controls) were trained to perform an optogenetic detection task. The task consisted in a succession of trials. First, the trial started with a no-lick period of 4 ± 1 seconds, which was reset by any lick in order to encourage the mouse to refrain from licking. Then, a blue light spot was projected on wS1 cortex through the intact optical window or on the dental cement with a 5.64 mW/mm2 light intensity. It consisted of a filled circle of diameter 300 µm (∼single barrel size). The spot was projected during 500 ms with a 50% duty cycle. We used the same DLP system as for the electrophysiological recordings. Once the spot was projected, the mouse had 2 seconds to lick a sensor to trigger a 20 µL water reward. Both spots (on the cortex and on the cement) were rewarded. To distinguish reflex lick from intentional lick, we used a threshold of 2 licks for the mouse to validate a trial and being rewarded (Ceballo et al., 2019a). Finally, the trial ended with an intertrial of 7 ± 1 seconds so that the mouse could consume the reward and get ready for the next trial. During the entire session, a blue masking light was placed in front of the mouse and directed towards its eyes to mask optogenetic projections.

Before training, all mice were first habituated to water regulation for 2 days. Then, mice were habituated to head-fixation and learned to lick a spout to obtain water during 2 sessions (one per day). During these habituation sessions, each lick triggered a reward. Each subsequent daily session lasted 30 minutes, during which we presented a set of trials in a pseudo-random order (spot onto the cortex or cement). After nine days of training, to rule out a potential visual contribution to the detection of the spot, we replaced the cement spot by a blank stimulus (no projection at all) for the two control mice.

### Discrimination task of a dynamic optogenetic stimulation over wS1

Five Emx1-Cre mice and 2 Scnn1a-Cre mice were trained to perform an optogenetic discrimination task. We used the same DLP system as for the electrophysiological recordings to precisely flash a 2D light pattern rotating on wS1 cortex through the intact optical window. Stimulation patterns were a light bar (700 µm long, 150 µm wide) oriented at 360 different angles (every 1°) and rotating on a disk of diameter 1.5 mm. We previously demonstrated that transgenic mice expressing ChR2 can accurately track these patterns when projected on a somatotopic space, such as wS1 (Lassagne et al., 2022). The disk was centered on the C2 barrel location but the barrel itself was never illuminated to avoid over-stimulation (see ***Fig. 4D***, white circle at the center). In pilot experiments (not shown), Emx1-Cre x Ai32 mice were able to learn this task using a photostimulation intensity of ∼0.94 mW/mm2. We therefore used the same intensity for Emx1-Cre mice injected with viral particles. We wished to apply similar light intensity to the main target cells of Scnn1a-Cre mice. To this aim, using the Allen Brain Atlas, we first measured the depth of wS1 L2/3, which is the most stimulated layer in Emx1-Cre mice (Yizhar et al., 2011), and of wS1 L4, the most stimulated layer for Scnn1a-Cre mice. We obtained depths of 150 µm and 430 µm. We estimated the relative proportion of blue light reaching these depths to be around 24% and 4% respectively (Yizhar et al., 2011). We thus applied a light intensity 6 times higher (5.64 mW/mm2) on Scnn1a-Cre mice. Details about light pattern sharpness and evoked spiking activity are available in (Lassagne et al., 2022).

Before training, all mice underwent the same habituation protocol as described above for the detection task. Each subsequent daily session lasted 30 minutes, during which we presented a set of trials in a random order. Each trial consisted in the presentation of one trajectory of the photostimulation bar, appearing at the most caudal position initially, and then rotating towards a rostral Rewardable zone (60° wide) with different kinetics. The bar position was updated every 10 ms with a 50% illumination duty cycle. Each trajectory was taken from a dataset of 252 pre-loaded trajectories, all lasting between 10 and 20 s and entering the Rewardable zone at least once (more details in (Lassagne et al., 2022)).

A lick had different consequences depending on the angular location of the photostimulation bar at that time. A lick when the light bar was in the Rewardable zone (green area, ***Fig. 4D***) led to an immediate 7 µl water reward. A lick in the No-lick zone (red area) immediately ended the trial and was followed by the 5 s intertrial interval, during which the cortex was not photostimulated. A new trial started immediately after. A lick in the Neutral zones (white areas) had no consequence. If the mouse drank more than 1.2 mL of water during one session, only one lick out of two or three was rewarded with water during the following sessions, starting with the first lick inside the Rewardable zone.

In the first experiments with Emx1-Cre mice, training was stopped after 10 days, when performance was already high (n = 2). Three more Emx1-Cre mice were trained for 15 days. As training duration varied across mice, the last session of each mouse was labelled “Last” for group analysis (10^th^ or 15^th^). Two Scnn1a-Cre mice were trained for 15 and 19 sessions. Emx1-Cre x Ai27 mice were trained for 5 days (n = 5), 7 days (n = 2) and 10 days (n = 1).

## Discussion

In this study, we have provided evidence for optogenetic control of cortical neurons *in vivo* using the PhP.eB serotype as a vector to express the ChR2-H134R opsin. We have shown that mice injected with this viral preparation can learn to extract an information of position from patterned optogenetic stimulation of the somatosensory cortex, and use it to guide their behavior. Furthermore, by combining different expression strategies (transgenic mouse lines and viruses), we were able to directly stimulate either layer 4 only, all other layers of the cortex, or all layers at the same time, during a discrimination task of dynamic cortical patterns. Learning occurred in all groups of mice but was much faster when we stimulated simultaneously excitatory neurons in all the layers.

### Benefits and limitations of the PHP.eB serotype for neuronal transgene expression in the cortex

To our knowledge, this study is the first to report the use of the PHP.eB serotype as a vector to express an excitatory opsin in the brain. Specifically, it allowed us to achieve widespread expression for mesoscale photoactivation *in vivo* of the cerebral cortex (***Fig. 3,4***). Indeed, the homogeneity of expression within a specific structure and/or a specific layer, obtained with the PHP.eB serotype in combination with a Cre driver mouse line, is a major advantage over intracerebral viral injections. For example, Grodem and colleagues (Grødem et al., 2023) reported a homogeneous expression of jGCamP8s with the PHP.eB strategy, whereas an intracerebral injection yielded a 40% higher fluorescence level at the center of the expression zone compared to its edges. Homogeneity is particularly important for patterned optogenetic applications. In our experiments, the projecting device delivers uniform light intensity across the field of projection. Coupled with homogeneous opsin expression, it guarantees the same level of activation for all illuminated zones in a given pattern.

Yet, the expression level and the proportion of transfected neurons are still limiting factors (Chakrabarty et al., 2013; Chan et al., 2017; Goertsen et al., 2022; Jang et al., 2023; Mathiesen et al., 2020). Our results show a transfection of all neocortical areas (***Fig. 1***), but we noticed an overall lower fluorescence in mice injected with the PHP.eB viruses compared to double transgenic animals. For example, the raw fluorescence from L4 neuropil was weaker in Scnn1a-Cre mice injected with PHP.eB particles than in Scnn1a-Cre x Ai32 mice (***Fig. 2***). Chan and colleagues (Chan et al., 2017) succeeded to transfect 51% of cortical DAPI+ cells (69 % of cortical neurons) using a dose of PHP.eB viruses similar to our own, yet without any specific promoter as Emx1. Another study (Mathiesen et al., 2020) reported a lower transfection rate with the same dose: 12% of DAPI+ cells transfected in the cortex, among which 83% were neurons. Apart from the variability, all these studies converge on the following observation: only a fraction of targetable cells are transfected. In comparison, studies using equivalent transgenic lines reported recombination in approximately 90% of all cortical neurons, suggesting a complete coverage of cortical excitatory neurons (Gorski et al., 2002; Guo et al., 2000; Madisen et al., 2012) .

### Absence of transgene expression in layer 4 barrels of S1

Unexpectedly, we found no or very low fluorescence in the L4 barrels of Emx1-Cre mice injected with the PHP.eB particles. (***Fig. 1B-D***). Because we observed a strong fluorescence in L4 barrels of double transgenic mice expressing the same construct (Emx1-Cre x Ai32, ***Fig. 1EF***), we hypothesized that this drop of fluorescence was probably due to the particular strategy of PHP.eB viral injections in adult mice of the Emx1-Cre driver line, more than to the gene construct itself.

More precisely, histological observations led us to conclude that L4 excitatory neurons (mainly stellate cells) of Emx1-Cre x Ai32 mice expressed the opsin in their neurites, known to be dense in the barrel neuropil, whereas the L4 neurons of Emx1-Cre mice injected with viral particles did not. We will discuss two potential explanations for this difference: capsid tropism and promoter strength changes across time.

### PhP.eB tropism

The viral capsid is a major determinant of tissue and cell type tropism (Chakrabarty et al., 2013). The PHP.eB serotype was engineered from the AAV9, for which a drop of fluorescence in L4 was already reported, in particular after intracerebral injections in cats and mice (Liu et al., 2020). For PHP.eB, a strong tropism has been observed for astrocytes and neurons, in particular in the cortex for inhibitory neurons (Brown et al., 2021; Jang et al., 2023) and L5 excitatory neurons (Brown et al., 2021; Grødem et al., 2023; Jang et al., 2023; Mathiesen et al., 2020; Michelson et al., 2019) . Jang and colleagues measured a slight negative bias for L4 excitatory neurons. Thus, the PHP.eB serotype could tend to favour other layers than L4, and this may have contributed to our observations (***Fig. Suppl. 2***).

Moreover, capsid tropism is highly dependent on the animal’s age at the injection time. In the days following mouse birth, the tropism of most AAVs is switching away from neurons towards astrocytes (Chakrabarty et al., 2013). In our study, mice were injected with viruses between 5 and 11 weeks old, probably reducing the fraction of transfected neurons.

### Emx1 expression in the cortex of the adult mouse

The Emx1 promoter is particularly strong during embryonic development, specifically in progenitors for cortical excitatory neurons (Gorski et al., 2002). Its activity then decreases rapidly in post-natal life (Gulisano et al., 1996). In the adult, the precise pattern of Emx1 strength across the different layers of the cortex is difficult to estimate, as most studies have focused on juvenile animals, and not necessarily on the somatosensory cortex (Chan, 2001; Gorski et al., 2002; Gulisano et al., 1996; Jin et al., 2000) .

In the double transgenic Emx1-Cre x Ai32 mice, the strong Emx1 promoter in the embryo results in a high level of Cre expression in neuronal progenitors. Thus, very early on, the Cre protein recombines lox sites of the ChR2 gene construct, and the defloxed ChR2 gene is passed on to daughter neuronal cells. By contrast, when we introduce a floxed gene construct via viral injection in Emx1-Cre mice at an adult stage (5-11 weeks old), i.e. at a time when the Emx1 promoter is probably much less active, recombination probably happens in a smaller fraction of neurons, and may depend on their laminar position. Indeed, we cannot exclude that the Emx1 promoter would be particularly weak in L4 of the barrel cortex in adult mice. In this respect, it is interesting to note that low expression of the transgene was also observed in Emx1-Cre adult mice injected with a virus containing a floxed calcium reporter (***Fig. Suppl. 2***). Overall, we conclude that the combination of PHP.eB capsid tropism and Emx1 promoter strength at the time of the viral infection, here in the adult mouse, is likely to be responsible for the particular laminar pattern of expression obtained in the “Emx1-Cre + virus” group of mice.

### Impact of the laminar pattern of optogenetic activation on the discrimination task

We verified first that using the PHP.eB strategy, we could elicit spiking activity in vivo and train mice to detect the neuronal activation of S1 induced by optogenetic spots (***Fig. 3***), similarly to what was achieved with a double transgenic strategy previously available (Abbasi et al., 2018; Ceballo et al., 2019b). We then trained mice to discriminate the location of a dynamic cortical elongated stimulus (***Fig. 4***). We measured striking differences in learning kinetics between the three expression profile groups (***Fig. 4H, Fig. Suppl. 6***). The number of training sessions necessary to reach the same level of performance was twice in the ‘all layers except L4’ and ‘only L4’ groups compared to ‘all layers’ group (***Fig. 4I***). Nonetheless, although the direct stimulation of excitatory neurons from all layers promoted faster learning, all groups of mice were able to learn the task and reached a similar high level of performance. These observations suggest that even if cortical stimulation was less easily discriminated at first in some groups of mice, cortical circuits were flexible enough to learn (Dalgleish et al., 2020; Houweling and Brecht, 2008; Luis-Islas et al., 2022). We conclude that the activation of layer 4 first was not essential to reach a high discrimination performance (***Fig. 4G-H, Fig. Suppl. 6***), but that it did speed up learning when all layers were stimulated simultaneously.

### Somatotopic map and receptive field size may influence perception

Beyond the global amount of depolarization or the number of activated neurons (Ceballo et al., 2019a), the location and the functional identity of the neurons activated by optogenetics could be important to explain these results. In previous work, we showed that learning to discriminate the position of a moving cortical stimulus (***Fig. 4***) was only possible when the stimulated area exhibited a continuous topographic map of the sensory periphery (Lassagne et al., 2022). This conclusion was drawn from optogenetic stimulation in mice with the “all layers” ChR2 expression profile. In these animals, L4 neurons, characterized by sharp receptive fields strongly centered on one whisker, were probably efficiently stimulated. This may have generated a perception corresponding to a precise subset of whiskers, albeit changing dynamically. By contrast in the ‘all layers except L4’ group of mice, L2/3 neurons were probably responsible for most of the perceptual effect, as L4 neurons receive very few inputs from all other layers of the column and were probably little activated (Lefort et al., 2009). L2/3 neurons are characterized by broad subthreshold receptive fields integrating inputs from several whiskers (Brecht et al., 2003; Le Cam et al., 2011; Sato et al., 2007). Thus, their activation could have generated a blurrier perception than when L4 neurons were directly activated. This difference could explain why animals of that group needed more sessions to estimate the position of the stimulus (***Fig. 4I***) and to lick in accordance with the position of the bar (***Fig. 4F-G***). However, pushing this hypothesis further, we could have expected that in the “only L4” group, direct co-activation of neurons with small receptive fields precisely organized on the somatotopic map could have favoured perceptual discrimination and rapid learning. Best efficiency of “only L4” stimulation could also have been expected given that L4 neurons are the first activated by peripheral stimuli, and should thus trigger a cascade of neuronal responses recapitulating activation of the well-described feedforward pathway receiving lemniscal inputs. The fact that learning was quite slow in the “only L4” mice suggests, however, that L4 direct activation was not, at first, successfully recruiting the cortical ensembles critical for perception. Perhaps one of the reasons was that the concomitant activation of cells in different layers of the same column is an important factor. Indeed, numerous studies pointed out that L5 pyramidal neurons integrate inputs across the column (Larkum et al., 1999) and that this integration correlates with perception (Takahashi et al., 2016).

### Conclusion and perspectives

In this study, we took advantage of the heterogeneous expression profiles obtained with the serotype PHP.eB to explore the behavioral effect of stimulating optogenetically different layers of the somatosensory cortex. Retro-orbital injection provides a way to express opsins in neurons in a minimally-invasive way. In the long term, this could be used to implement optogenetic artificial sensory feedback in closed-loop brain-machine interfaces (Abbasi et al., 2023, 2018; Goueytes et al., 2022). We found that directly activating all layers of the somatosensory cortex was the best strategy to train mice to discriminate cortical stimuli. In future experiments, it would be interesting to test the CAP.B10 serotype. Indeed, it has a higher neuronal tropism, transfects many more cortical excitatory neurons than PHP.eB in vivo, and transfects cortical layers more homogeneously (Brown et al., 2021; Jang et al., 2023). Furthermore, injecting the animals should be done at a younger age to improve the neuronal transfection. In this way, we might be able to improve the quality of the optogenetic feedback in order to restore an accurate artificial sense of touch.

## Acknowledgments

We thank Margot Peyré and Camille Gabot for help in training some of the mice in the discrimination task. We thank Henri Lassagne for preliminary experiments prior to this project. This work was funded by CNRS and Agence Nationale pour la Recherche (ANR PRC Motorsense, Hermin, Profound, Perbaco).

---

**Figure Suppl. 1.**
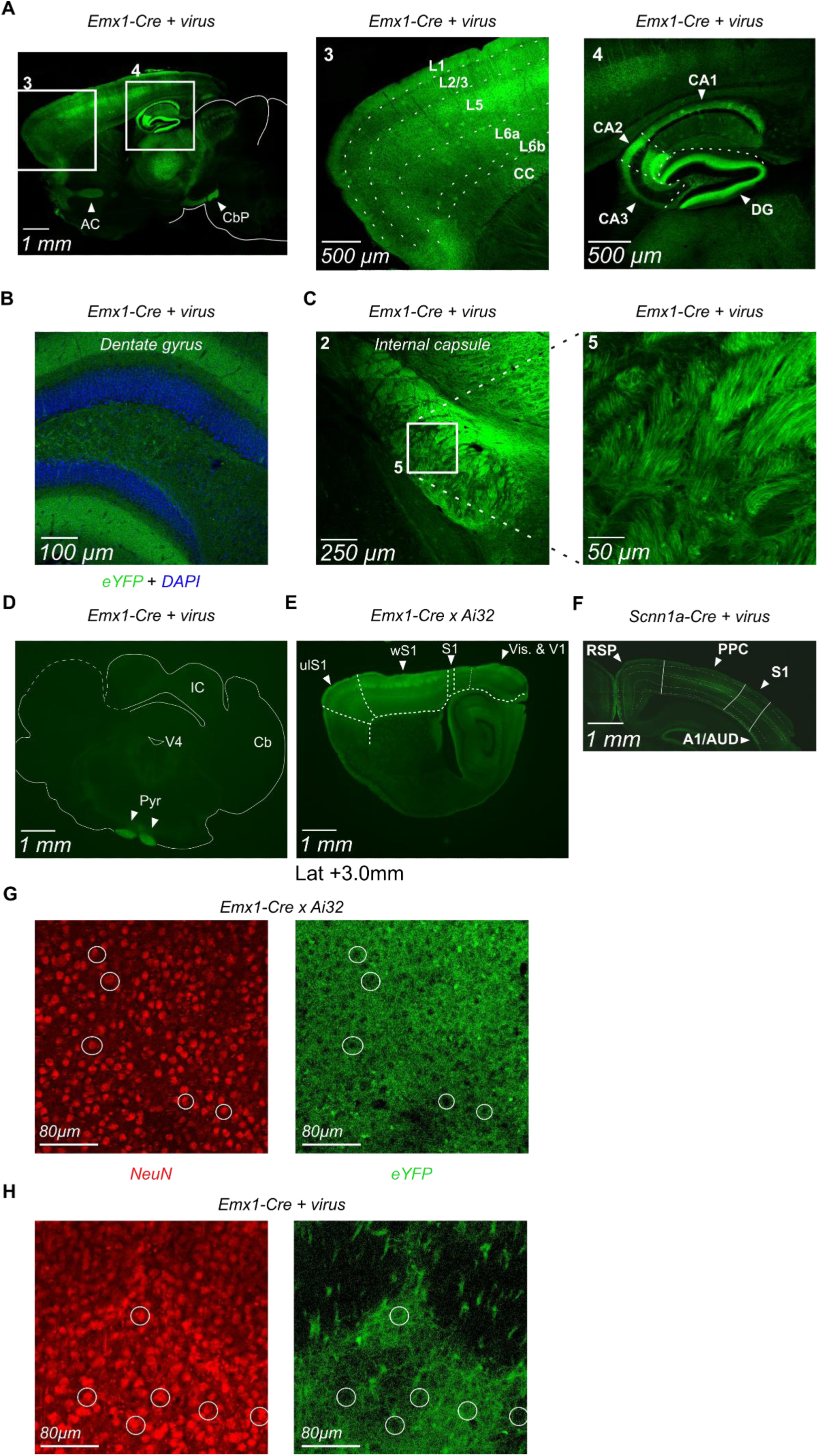
Transgene expression pattern in the brain of Emx1-Cre and Scnn1a-Cre mice after PhP.eB injection compared to equivalent transgenic mouse lines. **A Left** Fluorescence image of a sagittal section from an Emx1-Cre mouse brain showing expression of ChR2(H134R)-eYFP 4 weeks after AAV PHP.eB injection. The neocortex, hippocampus and white matter structures are regions of high fluorescence. The frontal pole (3) and the hippocampus (4) are outlined in white boxes. As expected with the Emx1 dependency, no fluorescence was observed in the cerebellum (delineated in white). Cerebellar peduncles display a high level of fluorescence as they convey cortico-cerebellar projections. AC: Anterior commissure; CbP: cerebellar peduncles. **Middle** Higher magnification of inset 3. Cortical layers are delimited, layer 5 displays the highest fluorescence level. L: layer; CC: corpus callosum. **Right** Higher magnification of inset 4. Among subregions of the Cornus Ammonis (CA), CA2 displays the highest fluorescence level. Fluorescence drops sharply at its boundaries. DG: Dentate gyrus. **B** Fluorescence image of the transgene superimposed with DAPI staining of a coronal section through the dentate gyrus from an Emx1-Cre mouse brain 4 weeks after AAV PHP.eB injection. Nuclei are densely packed in the granule cell layer and do not co-localize with the transgene. This illustrates that most of the fluorescence from the transgene comes from the neurites and that the opsin is expressed at the neuronal membrane. **C Left** Fluorescence image of a coronal section through the internal capsule from an Emx1-Cre mouse brain 12 weeks after AAV PHP.eB injection. Higher magnification of inset 2 from Fig. 1B. The internal capsule is a pure white matter structure (only axons). Several bundles of fluorescent axons are visible. **Right** Higher magnification of the inset 5 showing the axon-like morphology of the fluorescent fibers. **D** Fluorescence image of a coronal section from an Emx1-Cre mouse brain showing expression of ChR2(H134R)-eYFP 12 weeks after AAV PHP.eB injection. As expected, no fluorescence is observed from the midbrain, the hindbrain and the cerebellum due to the Emx1 dependency. Pyramids are fluorescent since they contain cortical axons projecting down to the medulla. IC: Inferior colliculus; Cb: Cerebellum; V4: fourth ventricle; Pyr: Pyramids. **E** Fluorescence image of a sagittal section from an Emx1-Cre x Ai32 mouse brain. In agreement with Fig. 1EF, the whole neocortex is fluorescent and the barrels are brighter than the neighboring layers. S1: primary somatosensory cortex; ulS1: upper limb S1; wS1: whisker S1; Vis. & V1: Visual areas and primary visual cortex. **F** Fluorescence image of a coronal section from a Scnn1a-Cre mouse brain 7 weeks after PHP.eB injection. Layer 4 of the PPC and auditory areas also show the highest fluorescence among their cortical layers. RSP: retro-splenial cortex; PPC: posterior parietal cortex; A1/AUD: primary auditory cortex and auditory areas. **G,H** Transgene fluorescence (green) and NeuN immunostaining (red) from a tangential cortical section including layer 4 of wS1 from an Emx1-Cre x Ai32 mouse brain (**G**) and from an Emx1-Cre mouse injected with PHP.eB (**H**). These images are higher magnifications of images in Fig. 1HI. Neuronal nuclei are circled (**Left**) and match with low-fluorescence areas of the eYFP pattern (**Right**), illustrating that the transgene is expressed at the neuronal membrane and not in the cytoplasm.

**Figure Suppl. 2.**
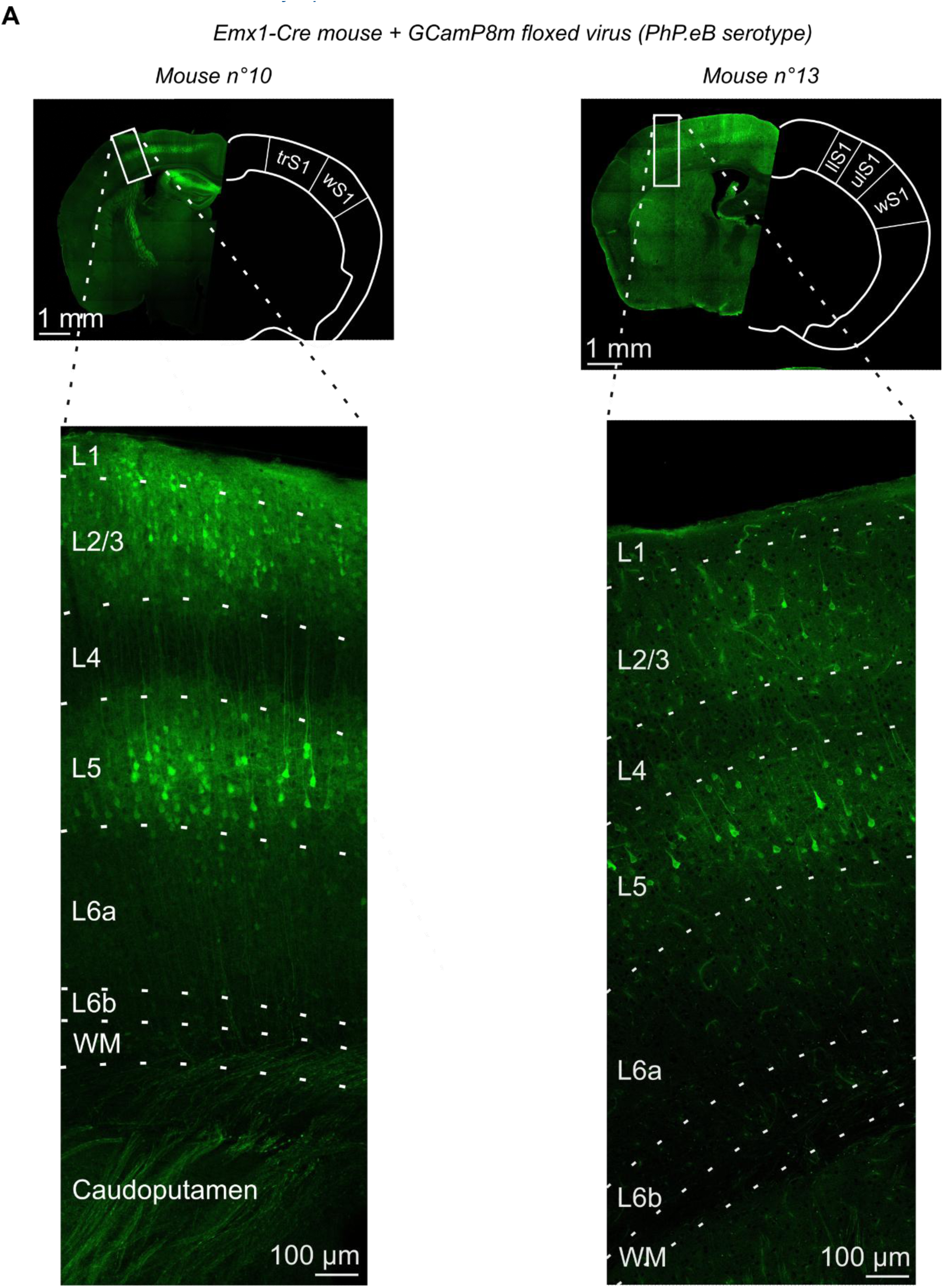
Expression of the cytoplasmic reporter GCamP8m via PHP.eB serotype also shows low fluorescence in layer 4 of Emx1-Cre mice. **A Top** Fluorescence image of a coronal section from two Emx1-Cre mouse brains showing expression of GCamP8m 4 weeks after AAV PHP.eB injection. Apart from the gene of interest (coding for GCamP8m and not ChR2-H134R) the construct is the same as before: Cre dependency and CAG promoter. Part of the whisker S1 area (wS1) is outlined a white box. llS1: lower limb S1; ulS1: upper limb S1; trS1: trunk S1. **Bottom** Higher magnification of the insets. Fluorescent somas are mostly located in layers 2/3 (L2/3) and layer 5 (L5). Neurons of layer 5 appear brighter than neurons of layer 2/3, probably because of their larger cytoplasm and consequently the higher amount of GCamP8m. There is almost no visible fluorescent soma in layer 4. The background noise and autofluorescence on the right image (mouse n°13) are probably due to the fact that the mouse has not been perfused.

**Figure Suppl. 3.**
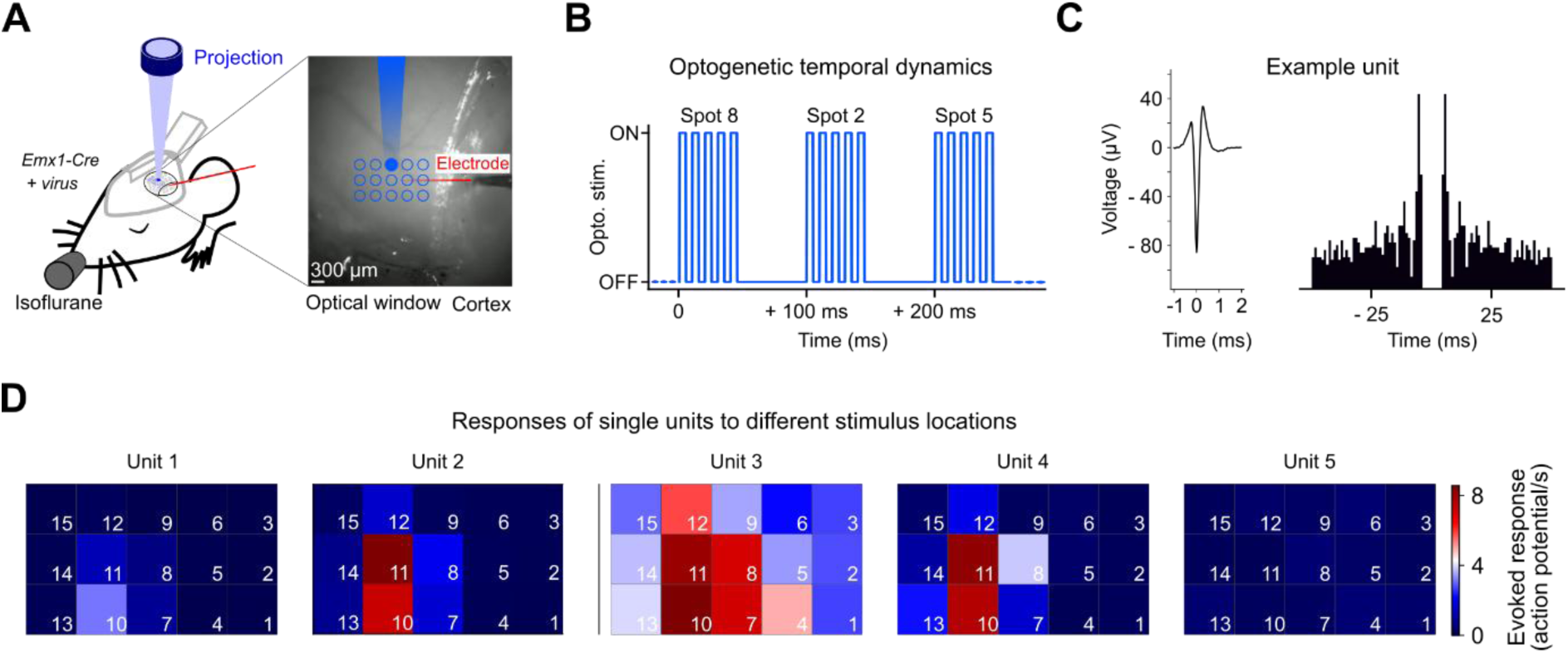
Spatial control of evoked activity in wS1 cortical neurons using patterned photostimulation in Emx1-Cre mouse expressing ChR2-H134R after AAV PHP.eB injection. **A Left,** Schematics of the setup. **Right**, An example spot (9^th^) is projected onto the window. Same as Fig. 3A. **B** Illumination pattern. A new spot was projected every 100 ms in a pseudo-random order. Each spot consisted of 5 successive flashes of 5 ms separated by 5 ms. **C** Example of the average waveform and autocorrelogram for one single unit after spike sorting and data curation. Single unit from Fig. 3C**-D**. **D** Response matrix for each unit, same quantification as Fig. 3D for the five sorted units. Average increase in the number of action potentials per second during the 80 ms following a light spot onset compared to the baseline period activity, for each of the 15 light spots.

**Figure Suppl. 4.**
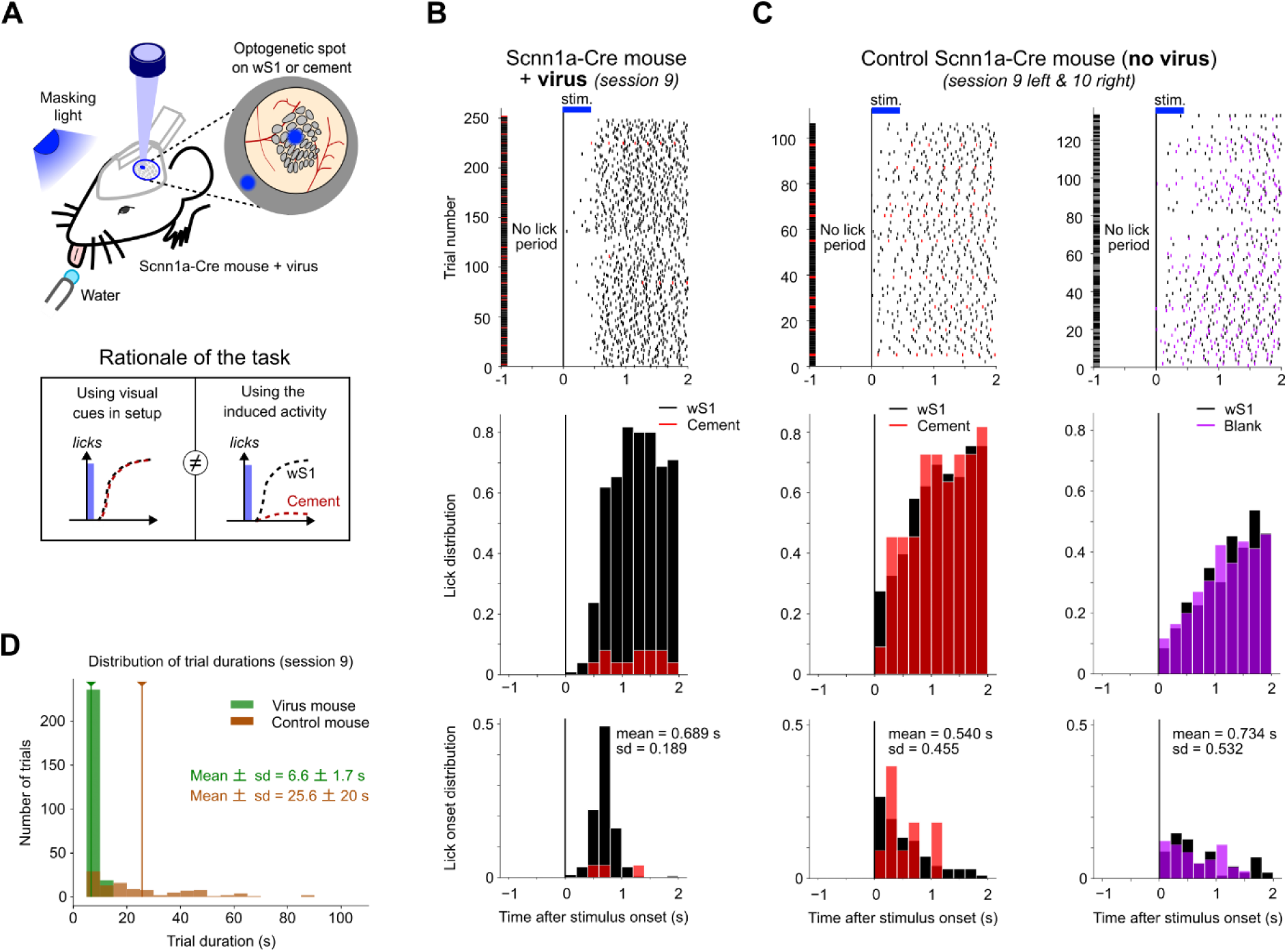
The detection task could be performed using either optogenetic cues or, if undetectable or absent, a timing strategy. **A Top,** Detection task, same as Fig. 3E**. Bottom**, Rationale of the task design. We wanted to figure out if the detection of the optogenetic flash was due to the activity it induced or by sensing it in any other way. By rewarding both spots, the mouse had every reason to lick as soon as it detected a spot, whether optogenetically or visually. Thus, the absence of licking for the spot on the cement indicates that it was not detected. **B** Licking activity for the Scnn1a-Cre mouse injected with the virus (“only L4”) during session 9 of the detection task. **Top, Middle** Same as Fig. 3G**. Bottom,** For each trial, we quantified the delay for ‘lick onset’ as the time of the first lick after the start of stimulation (reaction time). The mean lick onset distribution was computed over all trials of the session, same color convention as in Fig. 3G. The distribution of lick onset delays showed a single narrow peak (mean +/- s.d. = 689 +/- 189 ms). As the duration of the No Lick Period was random, this fixed reaction time to the stimulation further indicated that the mouse was indeed guided by the cue and not exploiting a timing strategy. **C Left,** Same representations as in **B** for a Scnn1a-Cre control mouse (no virus injected) during session 9 of the detection task. **Left Top**, Raster plots of licks. **Left Middle,** The lick distributions for both spots, through the window and on the cement, were similar, suggesting that there was no activation of neurons or retinal photoreceptors by light from the projector entering through the cranial window. **Left Bottom,** The distribution of lick onsets was not concentrated around one reaction time but as very broad (mean +/- s.d. = 540 +/- 455 ms), suggesting random initiation of licking bursts after the necessary still No-lick period. See also panel D. **Right,** Same analysis as **Left** for session 10 in the same control mouse, during which a spot was presented on wS1 or no stimulation was presented (no light or any cue at all) (purple curve and ticks). **Top and Middle,** In the absence of any stimulation, the animal still licked in the same way as it did for the spot on the cortex. **Bottom,** Reaction times to both spots were also similar. Overall, this suggests that the apparent detection of both spots by the control mouse was merely the effect of recurrent and sustained licking, i.e., a timing strategy that the mouse developed to take advantage of the temporal structure of the task. Due to technical issues, two consecutive partial sessions were run and pooled for each of sessions 9 (**Left**) and 10 (**Right**) of the control mouse. **D** Distributions of trial durations during session 9 for the virus-injected (green) and the control mice (orange). The duration of a trial was computed as the time spent in the No Lick Period (including all the resets) and the reaction period. The best theoretical mean trial duration (without any reset of the No Lick Period) would be 6 s. Arrows indicate the mean trial duration. Trial durations were much longer for the control mouse (up to 98 s, mean +/- s.d. = 25.6 +/-20.0 s) compared to the PhP.eB injected mouse (mean +/- s.d. = 6.6 +/- 1.7 s), mostly because of repeated resetting of the No Lick period. This difference confirmed the use of a timing strategy in the absence of opsin expression.

**Figure Suppl. 5.**
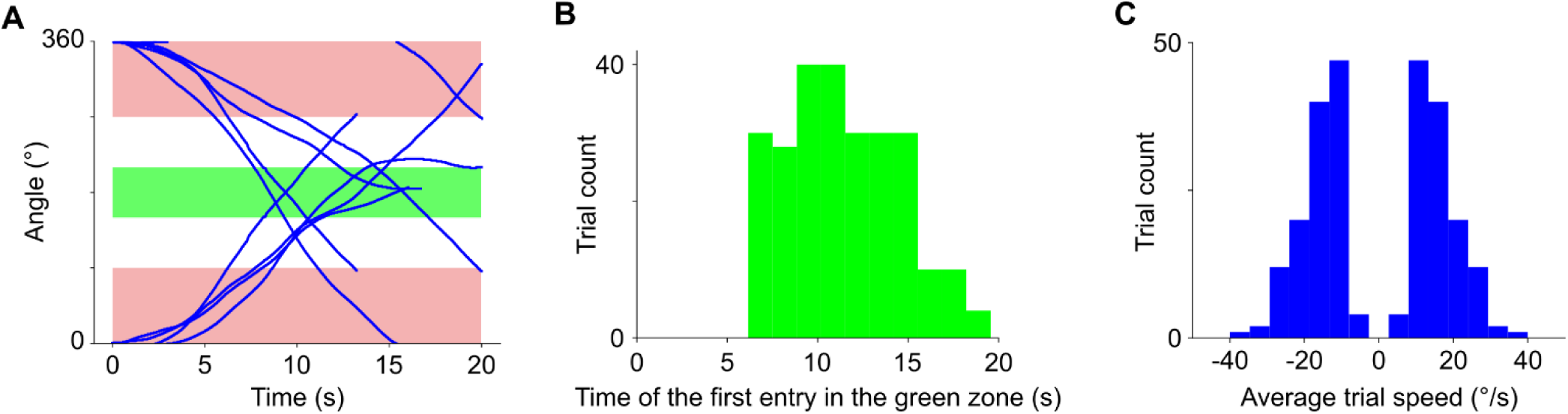
Dynamics of the optogenetic stimulus. **A** Eight example trajectories of the optogenetic stimulation. The reference angle 0° corresponds to the initial position on Fig. 4D, the most posterior one on the cortical surface. The photostimulation bar could rotate several times around the center, and could reverse directions. **B** Distribution of the times of first entry of the optogenetic stimulation in the Rewardable zone, for the 252 trajectories in the database. **C** Distribution of the average angular speeds of the optogenetic stimulation for the 252 trajectories in the database. The light bar can move either clockwise or counterclockwise.

**Figure Suppl. 6.**
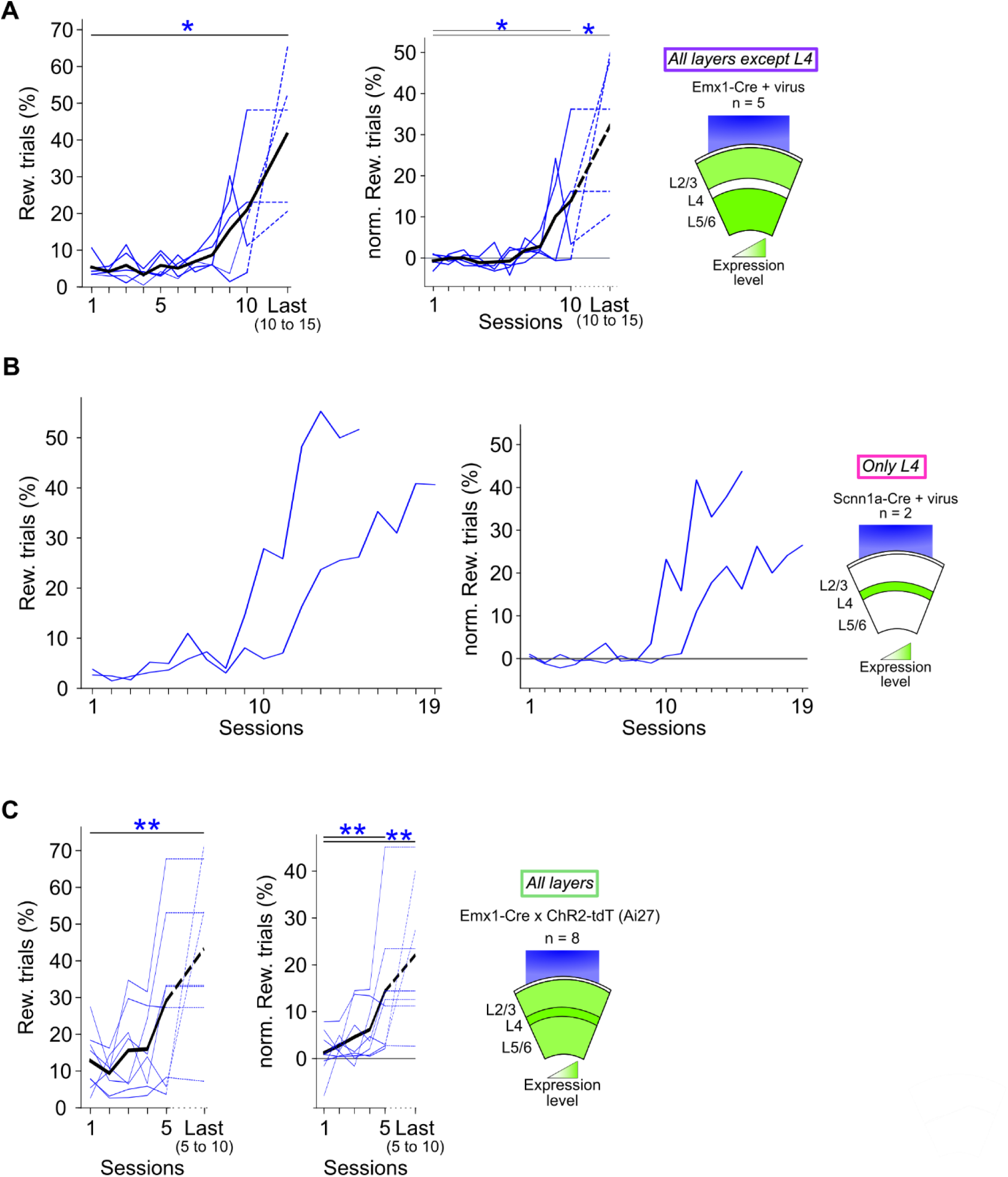
Raw and corrected learning curves of the three mouse groups in the rotating bar task. **A** Raw (Left) and normalized (Right) learning curves quantified by the percentage of rewarded trials for Emx1-Cre mice injected with PHP.eB (n = 5, **A**); Scnn1a-Cre mice injected with PHP.eB (n = 2, **B**); and Emx1-Cre x ChR2-tdT mice (n = 8, **C**). The normalized rewarded trial percentage is obtained by subtracting the chance level (see Methods). For the three mouse groups, the raw learning curves have similar shapes and dynamics to the corresponding normalized ones. The “Last” session differs between groups. For the group Emx1-Cre + virus, 2 mice were trained up to day 10 and three up to day 15. Scnn1a-Cre + virus mice were trained up to day 15 and 19. For the Emx1-Cre x Ai27 group, five mice were trained up to day 5, two mice up to day 7 and one up to day 10. Curves on the right are the same as Fig. 4H but truncated. Wilcoxon tests, * p < 0.05., ** p < 0.01.

